# Environmental stability of HPAIV H5N1 in raw milk, wastewater and on surfaces

**DOI:** 10.1101/2024.10.22.619662

**Authors:** Franziska Kaiser, Santiago Cardenas, Kwe Claude Yinda, Reshma K. Mukesh, Missiani Ochwoto, Shane Gallogly, Arthur Wickenhagen, Kyle Bibby, Emmie de Wit, Dylan Morris, James O. Lloyd-Smith, Vincent J. Munster

## Abstract

H5N1 influenza outbreaks in dairy cows necessitate studying potential transmission routes among livestock and to humans. We measured the stability of infectious H5N1 influenza virus in raw milk, wastewater, and on contaminated surfaces. We found relatively slow decay in milk, indicating that contaminated milk and fomites pose plausible transmission risks.

## Introduction

Since its first detection in cattle in March 2024, Highly Pathogenic Avian Influenza H5N1 (H5N1) virus has caused a multistate outbreak in dairy cows in the United States(1). The outbreak in dairy cows is caused by the Eurasian lineage goose/Guangdong H5 clade 2.3.4.4b that has been endemic in the USA since winter 2021-2022 (2, 3). H5N1 2.3.4.4b has caused a series of epizootic outbreaks in mammal species (3-5). During the ongoing US dairy cow epidemic, zoonotic and cross-species spillovers of H5N1 to farm workers and other mammalian species in proximity to dairy farms have occurred (6-8).

H5N1 virus replicates in mammary gland epithelial cells of dairy cows. Replication inside the udder results in high viral titers in milk from infected cows of up to 10^8.8^ 50% tissue culture infectious dose (TCID_50_) per milliliter (9, 10). Direct contact, environmental (via contaminated milk or wastewater streams), and mechanical transmission (via contaminated milking equipment) have been postulated as key drivers of cow-to-cow, zoonotic, and cross-species transmission (11, 12). To better understand environment and mechanical H5N1 transmission, we evaluated the stability of infectious H5N1 in raw milk, on surfaces, and in wastewater.

### The study

We assessed the decay rates and corresponding half-lives of H5N1 in raw milk, on polypropylene and stainless-steel surfaces, and in wastewater from a treatment plant. We spiked fresh, raw cow milk and wastewater with H5N1 influenza clade 2.3.4.4b isolated from the ongoing dairy cattle outbreak (A/bovine/OH/B24OSU-342/2024). We thank Richard Webby from St. Jude Children’s Research Hospital and Andrew Bowman from Ohio State University for providing the bovine H5N1 isolate. We tested samples of spiked fluids daily for seven days. To evaluate surface stability, we deposited four 12.5μl drops of spiked raw milk on stainless-steel or polypropylene disks. Samples were collected daily, by rinsing the disks with MEM tissue culture media. All experiments were performed in triplicate using biological replicates and quantified by endpoint titration on MDCK cells (Appendix).

We evaluated the stability in bulk milk and on surfaces at both 4°C and 22°C and the stability in wastewater at 22°C. We inferred posterior distributions for virus decay rates and half-lives using a Bayesian regression model (Figure 1 and Figure 2); below we report inferred values as the posterior median [95% credible interval].

**Figure 1:**
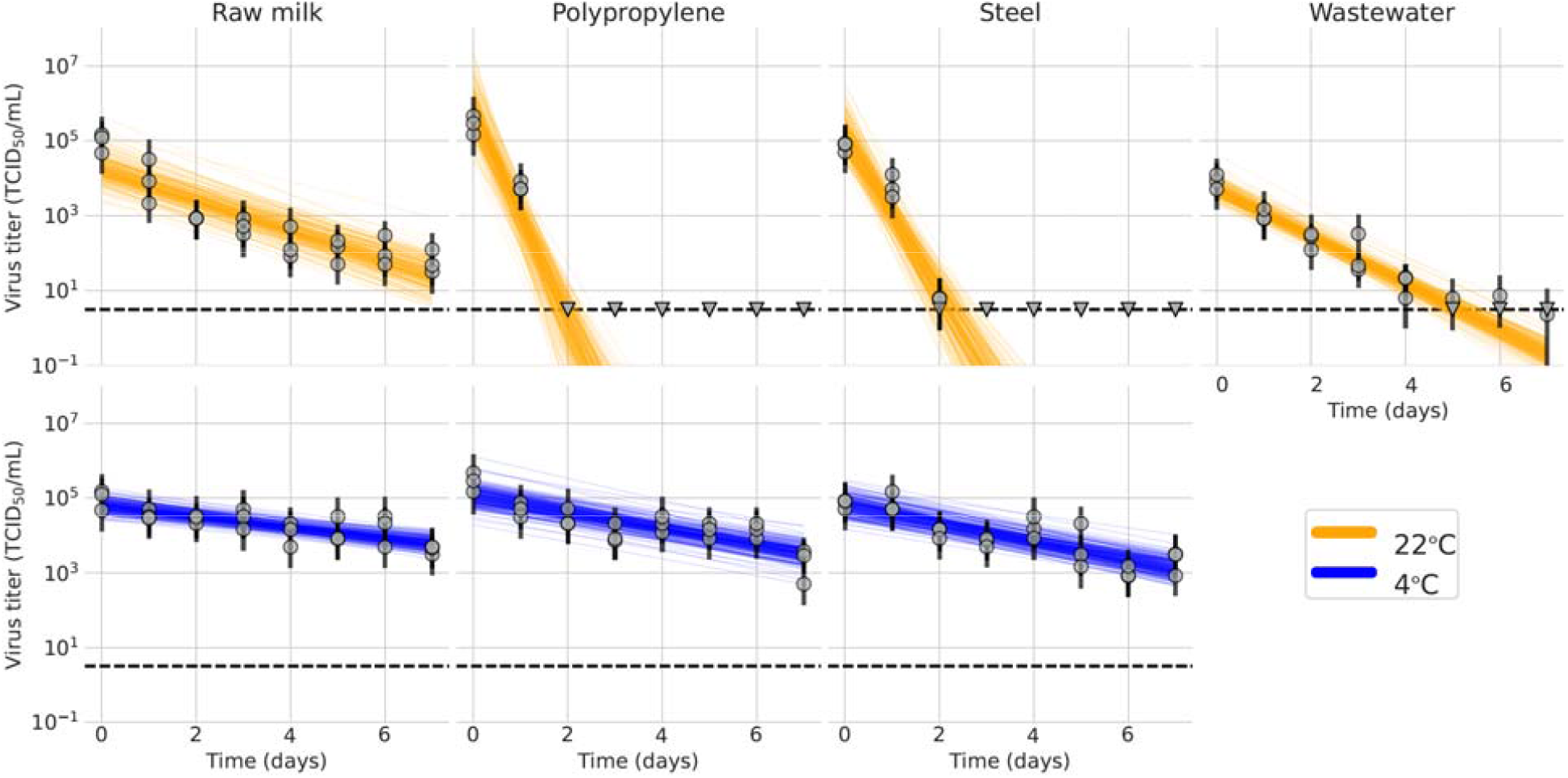
H5N1 stability in raw milk at 4°C (blue) and 22°C (orange), and in wastewater at 22°C (orange). The colored lines represent random draws from the joint posterior distribution of the exponential decay rate and the initial virus titer, where the intercept of each line is the initial titer, and the slope is the negative of the decay rate. The dashed horizontal lines are at 10^0.5^ 50% tissue-culture infectious doses (TCID_50_) per milliliter of medium and represent the approximate limit of detection.

**Figure 2:**
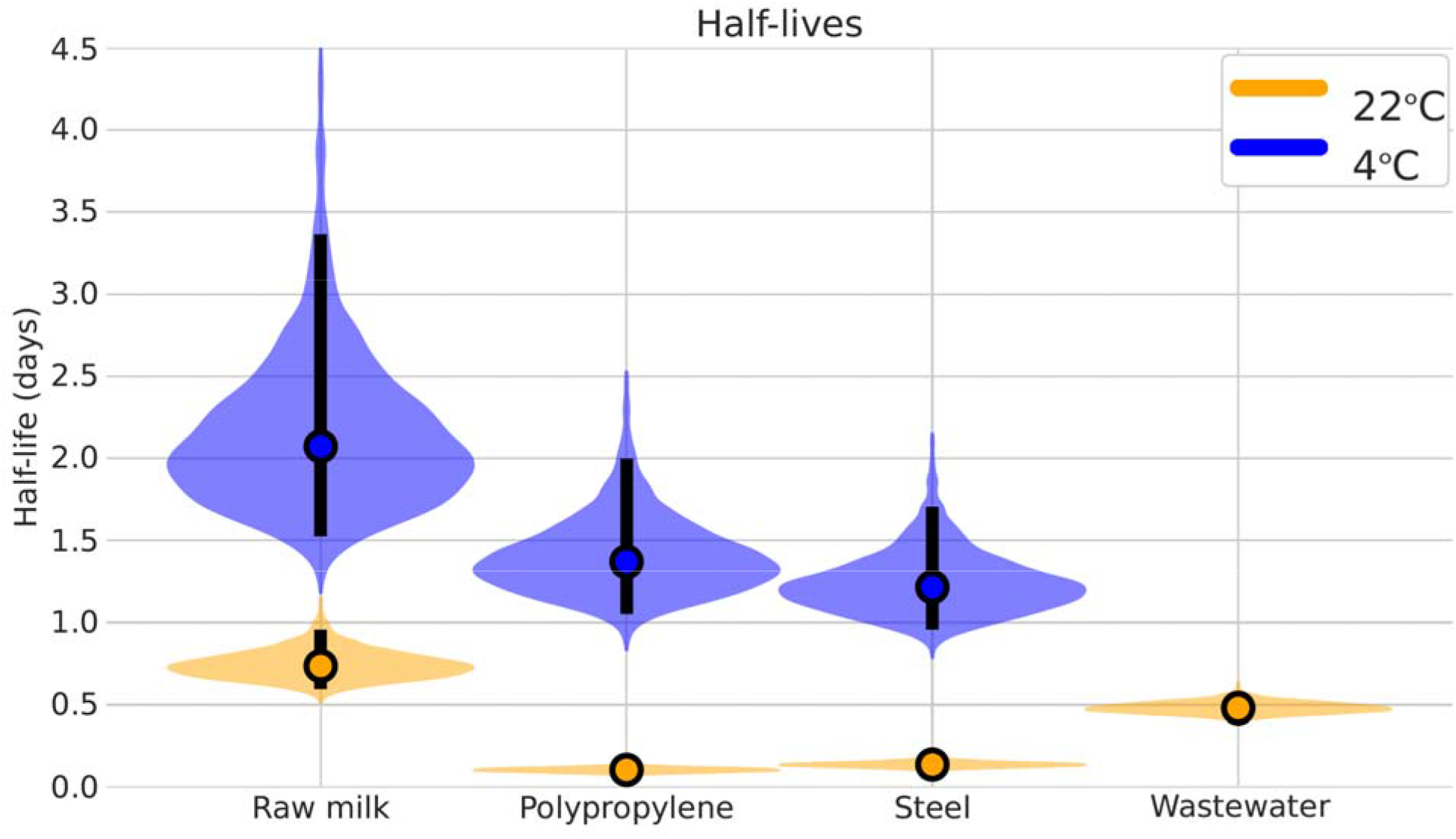
Calculated half-life of H5N1 in raw milk at 4°C (blue) and 22°C (orange), and in wastewater at 22°C (orange). Violin plots show the posterior distribution of the half-life of viable virus at each condition, determined from the estimated decay rates. The point at the center of each violin is the posterior median estimate and the black bar shows a 95% (2.5% – 97.5%) credible interval.

In bulk milk, we measured a half-life of 2.1 [1.5, 3.4] days at 4°C and 0.74 days [0.60, 0.96] at 22°C (Figure 2). The time needed for a 10 log_10_ reduction in virus titer was 69 [51, 112] days at 4°C and 24 [20, 32] days at 22°C. These results underline the potential of H5N1 to stay infectious in milk for multiple weeks, especially if refrigerated, given a sufficiently high initial titer.

We studied surface stability of H5N1 on stainless-steel or polypropylene surfaces at 4°C with 80% relative humidity and 22°C with 65% relative humidity. At 4°C, we measured a half-life of 1.4 [1.1, 2.1] days on polypropylene and 1.2 [1.0, 1.7] days on stainless steel. At 22°C, we measured half-life values of 0.11 [0.08, 0.14] days (2.5 [1.6, 3.4] hours) on polypropylene and 0.14 [0.11, 0.18] days (3.3 [2.5, 4.3] hours) on stainless steel (Figure 2). Decay rates of infectious H5N1 were comparable on stainless-steel versus polypropylene surfaces but were approximately ten times faster at room temperature (22°C) compared to 4°C. 10 log_10_ reduction in infectious virus titer would be achieved at 4°C after 45 [35, 62] days on polypropylene and after 40 [32, 57] days on steel. In comparison, at 22°C the 10 log_10_ decrease would take 3.6 [2.7, 4.6] days on polypropylene and 4.7 [3.7, 5.9] days on stainless steel.

In bulk wastewater at 22°C, the half-life of infectious H5N1 was 0.48 [0.42, 0.56] days (Figure 2). 10 log_10_ reduction of infectious virus in wastewater would take 16 [14, 19] days if stored at 22°C.

## Conclusion

Here, we investigated the environmental persistence of infectious H5N1 influenza virus in bulk raw milk, on contaminated surfaces, and in wastewater. For the bulk milk and surface stability we tested two environmental conditions, 4°C and 22°C, representing refrigerator and room temperature settings. H5N1 demonstrated high stability with a monophasic, exponential decay pattern in all experiments. As expected, refrigeration led to slower virus inactivation (13). Inactivation was slower in milk than in wastewater, possibly due to stabilization arising from milk’s high protein content, similar to mpox virus, where increasing protein content has been reported to increase stability (14).

With the estimated inactivation rate and a high virus titer of 10^8^ TCID_50_/ml (detected during the current outbreak) (9), detectable quantities of infectious virus could theoretically persist in refrigerated raw milk for 45 [33, 73] days. On surfaces, we did not find significant differences between exposure to stainless steel versus polypropylene but detected a high dependency on temperature. The half-life in wastewater of 0.48 [0.42, 0.56] days (12 [10, 13] hours) demonstrates persistence on a timescale that may lead to exposure of humans or other animals when in contact with contaminated wastewater or surface water.

The relatively high stability of H5N1 reported here, combined with reports of high H5N1 titers in milk from infected cows, highlights the potential for virus transmission by contaminated milk or fomites. This is consistent with postulated cow-to-cow transmission during the milking process, and with exposure to infected cattle herds leading to infections in dairy workers and other animal species at affected dairy farms. Infection risk via wastewater has not been demonstrated, but our results indicate the need for caution in disposing of milk from infected cattle. In addition, the detection of H5N1 sequences in wastewater during weekly wastewater sampling in 10 urban areas throughout Texas suggests widespread presence of H5N1 genetic material in wastewater in states affected by the outbreak in dairy cattle (15).

Further research is needed to connect the current findings to transmission risk via different pathways, particularly investigations of the probability of infection arising from different doses and routes of exposure to H5N1 influenza.

## Acknowledgments

This work was supported by the Intramural Research Program of the National Institute of Allergy and Infectious Diseases of the National Institutes of Health. S.C. and J.O.L.-S. were supported by the National Science Foundation (DEB-2245631).

## Biographical Sketch

Dr. Kaiser is a postdoctoral visiting fellow in NIAIDs Laboratory of Virology (Virus Ecology Section). She is interested in transmission of emerging viruses and outbreak responses. Mr. Cardenas is a doctoral student in the Biomathematics PhD program at the University of California, Los Angeles. He is interested in the emergence and control of viral pathogens.

## Notes

### Competing Interest Statement

The authors have declared no competing interest.

