## supplemental information for "Environmental stability of HPAIV H5N1 in raw milk, wastewater and on surfaces"

Appendix

**Experimental methods**

**Biosafety and ethics.** This study was approved by the Institutional Biosafety Committee (IBC) and performed in high biocontainment (BSL4) at Rocky Mountain Laboratories (RML), NIAID, NIH. All sample processing and sample removal from high containment followed IBC-approved Standard Operating Protocols (SOPs).

**Viruses and cells.** The HPAI H5N1 virus strain A/bovine/OH/B24OSU-342/2024, was obtained from Richard Webby (St. Jude Children’s Research Hospital) and Andrew Bowman (Ohio State University) and grown and titrated on Madin-Darby canine kidney (MDCK) cells in Minimal Essential Media (MEM) (Sigma-Aldrich, St. Louis, MO) containing 4µg/ml trypsin, 2mM L-glutamine, 50U/mL penicillin, 50µg/ml streptomycin, 1% NEAA, 20mM HEPES (all Thermo Fisher Scientific, Waltham, MA). MDCK cells were maintained in MEM supplemented with 10% fetal bovine serum (FBS) (Wisent Inc., St. Bruno, Canada), 2mM L-glutamine, 50U/mL penicillin, 50µg/ml streptomycin, 1% non-essential amino acids (NEAA) and 20mM HEPES.

**Viral RNA detection.** qRT-PCR was performed on RNA samples extracted from swabs or tissues using QiaAmp Viral RNA or RNeasy kits, respectively (Qiagen, Germantown, MD, USA). Viral RNA was detected with a Luna one-step RT-qPCR kit (New England BioLabs) designed to amplify total viral RNA from the matrix protein. Primers and probe used to amplify the viral RNA were as follows: forward primer 5’- AAGACCAATCCTGTCACCTCTGA-3’; reverse primer 5’- CAAAGCGTCTACGCTGCAGTCC-3’; probe FAM-TTTGTGTTCACGCTCACCGTGCC-TAMRA. Dilutions of RNA standards were numerically determined by droplet digital PCR, run in parallel, and used to estimate viral RNA genome copy numbers.

**Virus titration.** Viruses were titrated by endpoint dilution in MDCK cells. MDCK cells were inoculated with tenfold serial dilutions of the environmental stability samples. One hour after inoculation, cells were washed twice with phosphate-buffered saline (PBS) and supplemented with infection medium (MEM supplemented with 4 µg/ml L-1-tosylamido-2-phenylethyl chloromethyl ketone [TPCK] trypsin, 2 mM L-glutamine, 50 U/mL penicillin, 50 µg/ml streptomycin, 1% NEAA and 20 mM HEPES). Three days after inoculation, the supernatants of infected cell cultures were tested for agglutination activity using turkey red blood cells as an indicator of infection of the cells.

**Experimental design for stability studies.** Wastewater collection: approximately 1 L of untreated primary influent was collected from a municipal wastewater treatment plant in northern Indiana, United States, receiving an average flow of 11 million gallons per day (MGD). Immediately after being collected, the sample was stored overnight at −80 °C and, shipped to the Rocky Mountain Laboratories (RML) overnight on ice. The sample was then stored at −80 °C at RML until the experiments described herein were begun(1). Raw cow milk was collected from a local dairy farm in Montana. Prior to the experiment, fresh, raw cow milk was irradiated with 2M Rad and wastewater at 8M Rad to ensure inactivation of potential bacterial contaminants. Milk and wastewater were spiked with 10^6^ TCID_50_/ml of HPAIV H5N1 A/bovine/OH/B24OSU-342/2024. The virus-matrix suspension was then aliquoted into individual 1.5ml closed tubes and stored at the respective condition (4°C or 22°C). To maintain the desired temperatures consistently, we kept the aliquots in an environmental chamber (MMM Group, https://www.mmm-medcenter.com) at 22°C and 65% relative humidity, and a refrigerator (Fisherbrand™ Isotemp Laboratory Refriderator) at 4°C and 80% relative humidity. Samples were collected on 0d, 1d, 2d, 3d, 4d, 5d, 6d, and 7d timepoints and stored at −80°C until time of titration. All experiments were performed in triplicate.

**Experimental design for virus stability on surfaces.** Surface stability was evaluated on plastic (polypropylene, ePlastics) and AISI 304 alloy stainless steel (Metal Remnants), chosen to represent plastic and metal surfaces common in the dairy industry. 50µl H5N1 virus-milk suspension (2.5x10^7^ TCID_50_/ml; 1.25x10^6^ per 50µl) were deposited in four drops (12.5µl each) on polypropylene or stainless-steel disks. Disks were placed on plastic plates and contained in HEPA-filtered boxes under controlled environmental conditions. Experiments were carried out at 22°C and 65% humidity and 4°C and 80% humidity in an environmental chamber or laboratory refrigerator. Deposited virus was recovered by rinsing with 1ml MEM infection media at predefined time points (0d, 1d, 2d, 3d, 4d, 5d, 6d, and 7d) and samples were stored at −80°C until time of titration. All experiments were performed in triplicate.

**Statistical analyses.** The statistical analyses here closely follow those in Kaiser et al. 2024. As in that manuscript, we used a Bayesian approach to infer individual sample titers and virus half-lives from raw endpoint titration well data. We used Numpyro and Python 3 to specify and fit models. The principal difference between the analysis here and that in Kaiser et al. 2024 (2) is the inclusion of an additional Normal error term for predicted titers during environmental decay.

As previously described, we model well positivity in titration data with a Poisson single hit model; the hit probability depends on the initial virion concentration and the per-virion cell infection probability. In other words, if susceptible cells in a well are inoculated with infectious virions, the number of virions that infect cells and replicate is Poisson-distributed. However, only one virion needs to successfully enter a cell and replicate to produce a positive infection result for the well, so we model the well positivity as the probability that the Poisson random variable is greater than zero. The mean is measured in units of 50% tissue infectious dose (TCID_50_), so we do not need to consider the per-virion cell infection probability. We model the decay of viable virus as a linear decay in log_10_ units (as in Kaiser et al. 2024 (2)); however, to account for possible overdispersion in sample collection, we add an error term. For each sample *k* in experimental condition *j*, the quantity of viable virus at the time of sampling
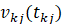
 is given by:

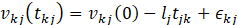

where
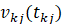
 is measured in log_10_ TCID_50_,
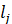
 is the rate of decay under condition *j*, and
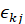
 is a Normally distributed error term with mean 0 and an inferred condition-specific standard deviation
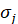
. The initial virus concentration
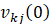
 is Normally distributed with inferred experimental condition-specific means
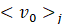
 and standard deviations
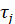
:

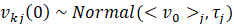
.

With this hierarchical approach, we model errors in the initial viral titer separately from errors in the collection and measurement of the titers during the decay process.

Prior distributions

In the following section, Normal distributions are parameterized as Normal (mean, standard deviation) and Positive Truncated Normal distributions (Normal distributions truncated to have only positive support) are parameterized as PosNormal (mode, standard deviation) where the mode and standard deviation are those of the underlying Normal distribution.

We used the following prior distributions:

For the individual titers
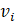
, we used a weakly informative Normal prior in units of log_10_ TCID_50_ per mL:

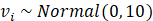

For the overdispersion of the titers, we placed a weakly informative PosNormal prior on the standard deviation of the error term
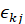
:

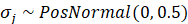

For the experiment-specific mean initial log_10_ titers
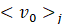
, we used a weakly informative Normal prior based on the target stock titer:

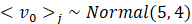

For the standard deviation of the individual sample initial titers
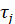
, we used a weakly informative Positive Truncated Normal distribution:

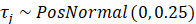

For inferring the half-life of infectious virus in days,
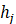
, we placed a weakly informative Normal prior distribution on the natural log of the half-life:

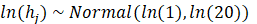

**Code and data availability**

All code and data needed to reproduce our analyses is archived on figshare (10.6084/m9.figshare.27277800), Github (<https://github.com/scardenas-0/h5n1-stability.git>) and Zenodo (https://doi.org/10.5281/zenodo.13947868), and licensed for reuse, with appropriate attribution and citation.
